# Assessing neuronal correlates of salience and their adaptability with naturalistic textures in macaque V1 and V2

**DOI:** 10.1101/2024.08.26.609744

**Authors:** Aida Davila, Adam Kohn

## Abstract

Salience is critical to vision. It allows stimuli that are different from their surroundings to ‘pop out’, drawing our attention. Perceptual salience is postulated to be encoded via a saliency map, based on differences in neuronal responsivity to simple image features at different spatial locations. Simple image features such as luminance, orientation and color are known to affect saliency and many of these features are encoded in primary visual cortex (V1), which several influential theories propose instantiate a saliency map. However, the degree to which more complex image features can determine salience, and whether there are neural correlates of salience which are computed outside of V1, remains unclear. Here we use displays of naturalistic textures to test for neural correlates of salience—termed pop-out responses—in V1 and area V2 of anesthetized macaque monkeys. Sensitivity to higher-order texture statistics arises in V2, so pop-out responses for these displays, if they exist, would be expected to be computed after V1. We presented displays in which a target texture, presented within the neuronal receptive field, was surrounded by distractors. Distractors could differ from the target texture in either higher-order texture statistics only, or in both lower- and higher-order statistics. We found little evidence for pop-out signals in either V1 or V2, for either display type. However, brief periods of adaptation could induce pop-out responses in V2. This suggests that adaptation might define which features of the environment are most salient, even if those features would otherwise not evoke pop-out responses.

**Significance statement:** We tested for neuronal correlates of salience—termed pop-out responses—using displays with multiple patches of naturalistic textures, whose higher-order statistics are encoded by neurons outside primary visual cortex. We found little evidence for pop-out responses in V1 or V2. However, brief periods of adaptation could induce pop-out responses in V2. Our results indicate that the computations that define bottom-up attention (i.e., salience) are malleable and continuously updated by our stimulus history.

---

Salience—the perceptual quality of a feature standing out from its surroundings—is determined in part by bottom-up (“pre-attentive”) mechanisms. A salience map has been proposed to guide our attention to the most salient location (Li, 1999; Itti and Koch, 2001). Features which elicit a stronger neuronal response in comparison to competitors are thought to determine to which feature and at what location bottom-up attention is drawn (Koch and Ullman, 1985; Treisman and Gormican, 1988; Li, 1999; Itti and Koch, 2001).

Computational (Koch and Ullman, 1985; Li, 1999; Itti and Koch, 2001), psychophysical (Treisman and Gelade, 1980; Bergen and Julesz, 1983; Wolfe, 1994) and physiological research has elucidated which features need to differ between a target and distractors for that target to be considered salient, and where within the visual cortex the relevant signals might be computed (Nothdurft et al., 1999; Hegdé and Felleman, 2003; Smith et al., 2007; Burrows and Moore, 2009; Poltoratski et al., 2017; Yan et al., 2018; Thayer and Sprague, 2023).

Primary visual cortex (V1) is thought to contain saliency maps that are computed on singleton features such as color, orientation, and shapes (Li, 1999). Surround suppression is a key mechanism for defining this map. Surround suppression refers to the ability of stimuli presented outside a neuron’s spatial receptive field to suppress the response to a stimulus within the receptive field (Hubel and Wiesel, 1965; Sillito and Jones, 1996; Sceniak et al., 1999; Walker et al., 1999; Angelucci et al., 2002; Cavanaugh et al., 2002a, 2002b; Angelucci and Bressloff, 2006; Carandini and Heeger, 2012). The tuning of this suppression is similar to that of the receptive field (Knierim and van Essen, 1992; Sillito and Jones, 1996; Cavanaugh et al., 2002b; Henry et al., 2013; Coen-Cagli et al., 2015). As a result, a target that differs from its homogeneous surroundings—a salient stimulus—will generate a stronger response in the relevant neurons than the responses of neurons encoding the surrounding stimuli (Li, 1999; Zhaoping and Zhe, 2015). A difference in a neuron’s responsivity for a target in a homogenous versus heterogenous displays—termed pop-out modulation—is often interpreted as a neural correlate of salience (Knierim and van Essen, 1992; Nothdurft et al., 1999; Lee et al., 2002; Hegdé and Felleman, 2003; Smith et al., 2007; Burrows and Moore, 2009; Dutta et al., 2016).

Though pop-out modulation is evident in V1, some have argued that this modulation is related to figure ground segregation rather than salience (Hegdé and Felleman, 2003; Burrows and Moore, 2009). V1 neurons also do not differentiate between salient and non-salient stimuli in displays constructed of conjunctions or singleton features (Hegdé and Felleman, 2003; Burrows and Moore, 2009), suggesting that salience for more complex displays may be defined in part by responses in higher visual areas. Consistent with this suggestion, pop-out modulation has been reported in V2 (Lee et al., 2002) and V4 (Mazer and Gallant, 2003; Burrows and Moore, 2009). However, whether this modulation arises de novo in higher areas or is inherited from V1 is unknown.

In this study we sought to test whether pop-out modulation occurs for displays in which target and distractor distinctiveness is defined by features encoded after V1, by measuring pop-out modulation for displays of naturalistic textures in macaque V1 and V2. Neurons in these two areas have distinct selectivity for textures (Freeman et al., 2013; Ziemba et al., 2016, 2018, 2019). V1 neurons are insensitive to high-level features of a texture, responding similarly to naturalistic textures and their spectrally-matched noise counterparts (Freeman et al., 2013; Ziemba et al., 2016). In contrast, V2 responds more strongly to naturalistic textures than to noise counterparts (Freeman et al., 2013; Ziemba et al., 2016, 2018). If pop-out modulation arises solely from mechanisms within V1, where neurons are blind to higher-order texture statistics, then pop-out modulation should be absent for displays which vary in these statistics, in V1 and areas downstream. On the other hand, if pop-out responses can be generated downstream of V1, they might be evident in V2 despite their absence in V1.

Our second objective was to test whether pop-out signals for texture displays could be altered by adaptation. Salience is not fixed. It can be altered by long-term learning (Julesz, 1981; Treisman and Gormican, 1988) and even by immediately prior sensory experience—or adaptation (McDermott et al., 2010; Wissig et al., 2013). Whether adaptation can alter pop-out modulation for displays of complex images is unknown.

We found little evidence for pop-out modulation for displays consisting of targets and distractors which contained distinct higher-order texture statistics, or even for displays in which targets and distractors differed in both spectral and higher-order statistics. However, adaptation could induce pop-out modulation, suggesting one function of adaptation may be to determine which features in the environment are most salient.

## Materials and Methods

### Surgery

All procedures were approved by the Institutional Animal Care and Use Committee of the Albert Einstein College of Medicine and were in compliance with the guidelines set forth in the National Institutes of Health’s *Guide for the Care and Use of Laboratory Animals*.

Recordings were conducted in four adult male monkeys (*macaca fascicularis*). Prior to the induction of anesthesia with ketamine (10 mg/kg), animals were administered glycopyrrolate (0.01 mg/kg) and diazepam (1.5 mg/kg). Once anesthesia was confirmed, animals were intubated and placed on 1.0%-2.5% isoflurane in a 98% O_2_ -2% CO_2_ mixture. Intravenous catheters were then inserted into the saphenous veins of both legs. Animals were placed in a stereotax. A craniotomy and durotomy were performed over V1, centered ∼5mm posterior to the lunate sulcus and ∼10mm lateral to the midline.

Postsurgical anesthesia was preserved with a venous infusion of sufentanil citrate (6–24 microg/kg/h, adjusted as needed) in Normosol solution with dextrose. To minimize eye movements, vecuronium bromide (150 microg/kg/h) was also administered via venous infusion. Vital signs, comprising of heart rate, SpO2, ECG, blood pressure, EEG, end-tidal CO2, core temperature, urinary output, and airway pressure, were continuously monitored to ensure suitable anesthesia and physiological state. Rectal temperature was maintained near 37°C using heating pads. Pupils were dilated with topical atropine. Gas-permeable contact lenses were used to protect the corneas. Additional supplementary lenses were used to bring the retinal image into focus. Antibiotics (Ceftliflex, 2.2 mg/kg) and a corticosteroid (dexamethasone, 1 mg/kg) were administered daily.

### Recordings

Neuropixel Phase 3B probes, placed on a custom 3D-printed probe holder, were lowered into the cortex using a microelectrode drive (Alpha Omega). Each recording session involved 2-4 Neuropixel probes, arranged mediolaterally. Neuropixel probes were sharpened prior to insertion; no guide tubes were used. Once Neuropixel probes were fully positioned in the brain, the craniotomy was filled with agar or Dura-Gel (Cambridge NeuroTech) to prevent desiccation of the cortical surface.

In one animal, neural responses were acquired from 384 channels extending over a distance of 3.84 mm, closest to the probe tip which was in V2. The remaining 3 sessions used a configuration which recorded from 384 channels in a staggered manner, spanning 7.68 mm of the probe and all layers of V1 and V2. Data was acquired with SpikeGLX software; signals were digitized at 30 KHz. Raw data were spike sorted with Kilosort 2.5, which clusters units based on waveform shape (Pachitariu et al., 2016). Kilosort output was then manually curated using Phy (https://github.com/cortex-lab/phy). Manual curation involved merging and splitting units and confirming waveform templates were from a single neuron.

### Visual stimuli

Stimuli were presented on a calibrated cathode ray tube monitor (Hewlett Packard p1230; 1024 × 768 pixels; 100 Hz frame rate, ∼40 cd/m^2^ mean luminance). The monitor was placed approximately 90-110 cm from the animal. Visual stimuli were generated and presented using custom OpenGL software (EXPO; P. Lennie). We first measured the spatial receptive field of each recording site by presenting small drifting gratings at 169 different screen locations, spanning a 5° × 5° region of visual space. Receptive field data were collected separately for stimuli presented to each eye. Recorded voltage traces were high pass filtered at 300 Hz and thresholded, and V1 and V2 aggregate receptive fields were computed from these multiunit responses. Target stimuli were centered on either the V1 or V2 neurons’ receptive fields on a single probe; in some experiments, stimuli were then re-centered over the receptive fields of neurons in the remaining area and additional data collected.

We presented textures synthesized with the Portilla and Simoncelli algorithm (2000). The algorithm quantifies a variety of statistics of an input texture and synthesizes new images enforcing the measured statistics. Texture synthesis for each image was initialized with a Gaussian white noise image. Analyzed statistics from the Portilla and Simoncelli algorithm were then iteratively applied (n = 50) so that the Gaussian white noise image resembled the original texture. Textures were synthesized with 4 scales and orientations of Gabor-like filters, and correlations were derived within a 15-pixel square.

Source textures were obtained from the normalized Brodatz texture database (Abdelmounaime and Dong-Chen, 2013), Gustaf Kylberg database (Kylberg, 2011), Oulu texture database (Lee and Bang, 2019), and Salzburg texture images (https://www.wavelab.at/sources/STex/). Each source texture (256 x 256 pixels) was gray scaled and served as the basis for a texture “family”. A total of 16 source textures were used. For each synthesized texture, we also synthesized a noise counterpart by randomizing the phase of the Fourier-transformed synthesized texture. All synthesized textures and noise images were adjusted to have the same contrast (RMS = 0.314) and mean luminance (128, on a 8-bit 0-255 range). Synthesized textures and noise counterparts were 512 × 512 pixels. We presented stimuli in a 4-degree aperture (40-54 pixels/degree), centered at random locations of the synthesized texture or noise stimuli. Each randomized centering of a texture family thus contained the same statistics (on average) but resulted in a distinct image. We refer to each randomized centering as a sample of the texture family.

To measure pop-out modulation, we presented displays consisting of either a target alone, target and distractors (which could be matched to the target stimulus or different from it), or distractors alone. We used displays of: (1) static grating stimuli, in which the target and mismatched distractors differed in orientation (and position) but were identical in all other aspects; (2) texture targets in which the distractors were noise images from the same texture family (or noise targets with texture distractors); (3) texture targets with other textures (i.e., from a different family) as distractors. For this final experiment, we chose randomly on each trial one texture to be the target and another to be the distractors from a set of two different naturalistic texture families. Each individual image—whether a target or distractor—was a different sample, at each location and on each trial. Data for experiment (3) were obtained from the unadapted or control trials, within the adaptation experiment described next.

In our adaptation experiments, trials consisted of a 600 ms adapter, a 100 ms blank, a 300 ms test period, and a 300 ms inter-trial interval. We adapted all target and distractor locations using the same Manhattan grid as the homogeneous/heterogeneous test condition. The adapter on each trial could be a uniform gray background (control), a set of one texture family, or a set of its noise image counterpart. For texture and noise adapters, the adapter could be from the same texture family as either the test target or distractors. Each texture or noise stimulus patch within our adaptation grid was a unique sample. Adapters consisted of a sequence of six randomly chosen samples, each displayed for 100 ms, to prevent retinal fading (Martinez-Conde et al., 2004). We did not present a new sample every 100 ms for the test stimuli (in these experiments or those described above) for fear that this might interfere with the generation of pop-out modulation. Each adaptation condition was presented 37-78 times (mean = 58).

### Analysis

Analysis was performed in MATLAB (MathWorks). Each neuron’s response was measured as the spike count in a 300 ms window during the test stimulus presentation. The window for both V1 and V2 units began 70 ms after stimulus onset and ended 70 ms after stimulus offset to account for response latency. Spontaneous activity was measured either during the blank interstimulus interval (100 ms) or, for the adaptation experiments, during the blank test stimulus which followed a blank adapter. We only included units in our analyses for which the mean response to at least one of the test stimuli in the target only condition was higher than the mean spontaneous response plus the standard deviation of that response. In addition, to ensure that the distractors did not encroach on the receptive field, we required units to have a response equal to or smaller than one standard deviation above the spontaneous activity, during the distractor only condition. Neurons were also required to have a median spike count of at least 1 spike/trial in the target only condition.

The pop-out index was calculated as the difference between the evoked response to a heterogeneous and homogeneous display, divided by the sum. We quantified the strength of surround suppression for each neuron through a suppression index, calculated by taking the difference between the evoked response for the homogenous display and the target alone condition and dividing by the sum.

To quantify the effects of adaptation, we calculated an adaptation index as the difference between the evoked adapted and unadapted response, divided by the sum. Units that met the inclusion criteria described above and were responsive to the test stimulus either before or after adaptation were included in the plotted distributions and statistical analysis.

We used bootstrap analysis to evaluate the statistical significance of the pop-out, suppression, and adaptation index for individual units. For the pop-out index, we pooled responses to homogeneous and heterogeneous displays and randomly chose subsets of responses (equal to the number of trials run in the original experiment) with replacement to create ‘homogeneous’ or ‘heterogeneous’ responses. We calculated the pop-out index for these data and then repeated the whole procedure 1,000 times. We defined the experimental value as statistically significant if it was in the top or bottom 2.5% of the bootstrapped distribution. Statistical significance of the other indices was computed similarly.

## Results

We used Neuropixels probes to record V1 and V2 neurons in 4 anesthetized macaque monkeys (Fig. 1A). In each session, we measured the spatial receptive fields at each recording site (see Methods). The location of a target stimulus was then centered on the aggregate receptive fields of either the sampled V1 or V2 units, in turn (Fig. 1B, plotted in green and blue, respectively).

**Fig 1.**
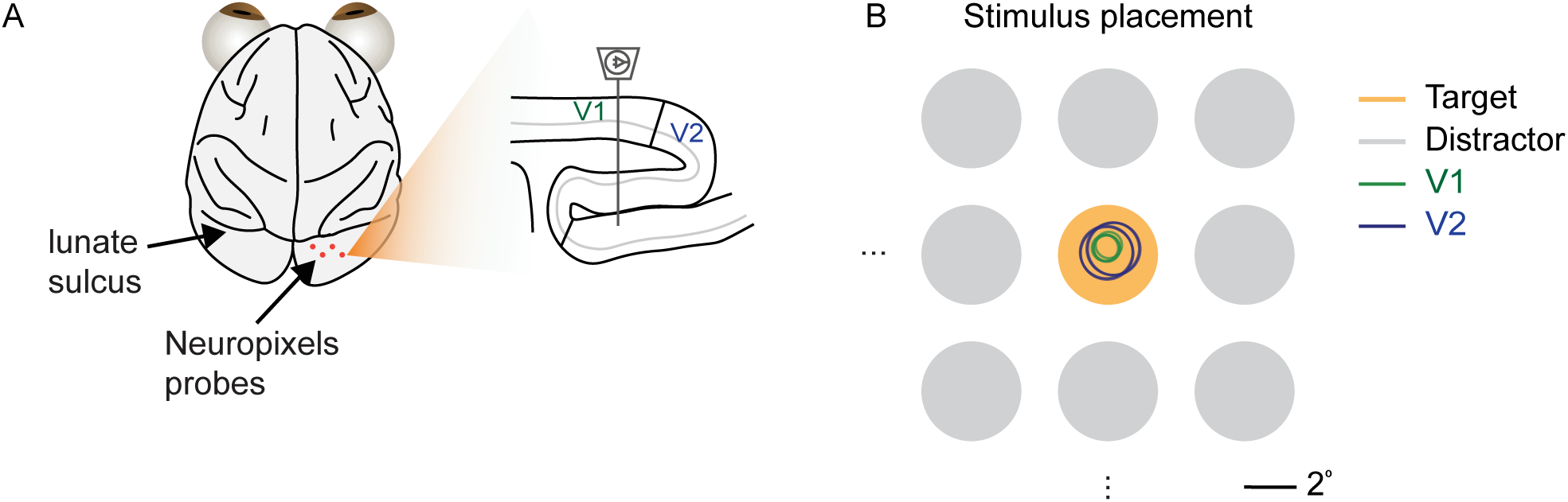
Recording approach. (A) Neuropixels probes (position on cortical surface indicated in orange) were lowered to record V1 and V2 units, shown in sagittal view (right). (B) Example receptive field locations for four V1 (green) and V2 (blue) units with respect to the center of the 4° target stimulus (orange). Distractors were spaced so as not to elicit a response when presented on their own.

To measure pop-out modulation and understand its basis, we used four types of displays (Fig 2A): (1) a target stimulus within the recorded units’ receptive fields, presented alone; (2) a heterogeneous display, defined as a target surround by distractors which differed from the target; (3) a homogeneous display, defined as a target surrounded by matched distractors; and (4) a display of distractors from the heterogenous condition, with no target stimulus.

**Fig 2.**
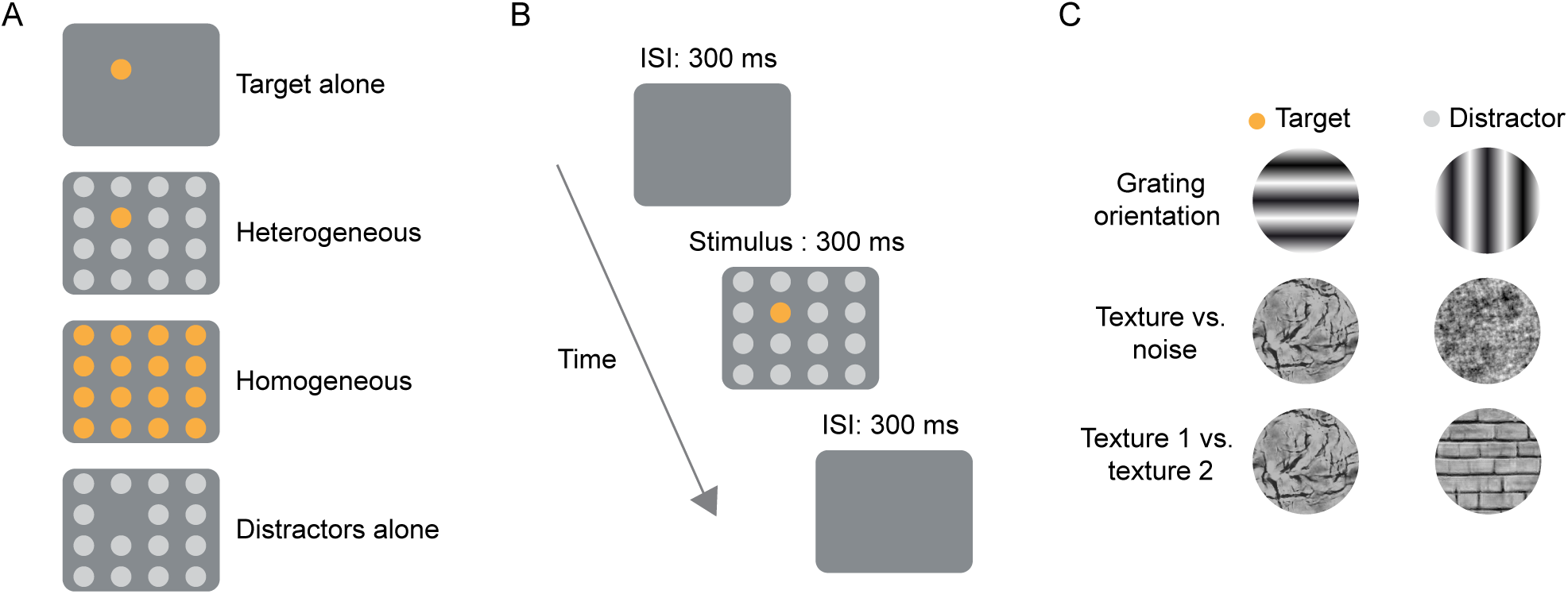
Visual displays. **(A)** We used 4 stimulus conditions: target alone, heterogeneous displays, homogenous displays, and distractors with no target. (B) Each trial consisted of a 300 ms presentation of a randomly chosen stimulus condition followed by a 300 ms interstimulus interval (ISI). **(C)** We used 3 different sets of target and distractor pairings.

Each stimulus patch was 4° in size and placed 1.2° (edge-to-edge) from its nearest neighbor, in a Manhattan grid arrangement. We used 4° patches because sensitivity to texture statistics involves modulation from the near surround (Ziemba et al., 2018) and because texture statistics involve dependencies across locations of an image which can be ill-defined for smaller images (Ziemba and Simoncelli, 2021). Distractors were placed in the units’ surround and did not elicit a measurable response. The number of distractors ranged from 19 to 29, depending on the position of the receptive fields on the screen. In all cases the number of distractors was chosen so that distractors extended to the edge of the visual display. One stimulus condition was chosen at random on each trial and presented for 300 ms (Fig 2B), followed by a 300 ms inter-stimulus interval (ISI).

In a given recording, distractors and targets could consist of either: (1) sinusoidal gratings of orthogonal orientation; (2) textures and their spectrally-matched noise images (described in further detail below); or (3) two different naturalistic textures (Fig 2C).

### Pop-out modulation for gratings

We first assessed pop-out modulation in V1 and V2 using static sinusoidal gratings. Targets and distractors were matched in luminance, contrast, and spatial frequency but differed by 90° in orientation (Fig 2C). We performed experiments with gratings to ensure that our choice of stimulus size and spacing evoked pop-out modulation, as reported in previous studies using bar stimuli (Knierim and van Essen, 1992; Hegdé and Felleman, 2003).

To quantify pop-out modulation, we used a pop-out index defined as the difference between the evoked response to a heterogeneous and homogeneous display, divided by their sum (Lee et al., 2002; Hegdé and Felleman, 2003; Smith et al., 2007; Burrows and Moore, 2009; Dutta et al., 2016). There was significant positive pop-out modulation in both V1 (mean pop-out index of 0.05±0.02, p=0.005 for difference from 0, t-test; Fig 3A, top) and V2 (0.12±0.01, p<0.001, t-test; Fig 3A, bottom), indicating stronger responses to heterogeneous displays. Pop-out indices were slightly higher in V2 than V1 (p=0.02, unpaired t-test). We also evaluated the percentage of individual neurons for which the pop-out index was statistically significant, using a bootstrap test (95% confidence interval; Fig. 3A, filled bars). More V2 units (39.1%) than V1 units (22.9%) had significant pop-out responses. Thus, grating targets and distractors differing in orientation result in clear pop-out responses in both V1 and V2 units.

**Fig 3.**
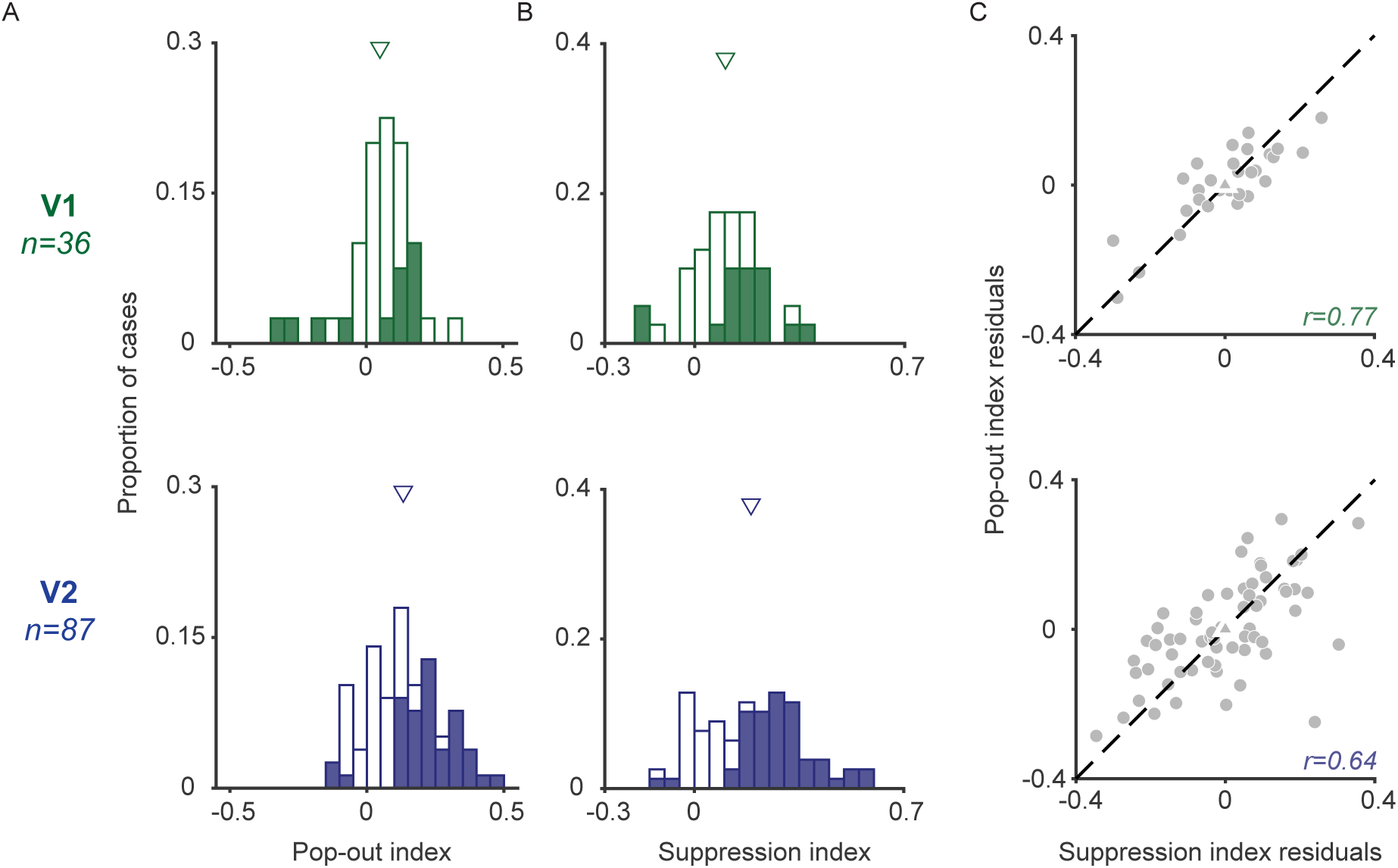
Pop-out modulation and suppression for displays of sinusoidal gratings. **(A)** Pop-out indices for V1 (top) and V2 (bottom) units. Shaded bars represent significant cases. Triangles above distribution indicate the mean. **(B)** Suppression indices for V1 (top) and V2 (bottom) units. Shaded bars represent significant cases. Mean is indicated by triangles above each histogram. **(C)** Relationship between suppression (abscissa) and pop-out (ordinate) index residuals, after regressing out the influence of responses to the homogeneous displays for each index, in V1 (top) and V2 (bottom).

Pop-out modulation is thought to arise from tuned surround suppression—the suppression of a neuron’s response to a stimulus in its receptive field by similar stimuli in the surround. We measured surround suppression using a suppression index, defined as the difference between evoked responses to the target alone and the homogeneous condition divided by their sum. A suppression index greater than 0 indicates weaker responses in the presence of distractors; an index less than 0 indicates a faciliatory influence of distractors. Neuronal responses in both visual areas were suppressed by distractors (mean suppression index in V1: 0.12±0.03, p<0.001, t-test; in V2: 0.21±0.02, p<0.001, t-test; Fig 3B). Suppression was stronger in V2 than V1 (p=0.006, unpaired t-test) and more V2 units (69%) had significant suppression indices than V1 units (57.1%).

We next asked whether units with stronger surround suppression were those for which there was stronger pop-out modulation. The two forms of modulation might be related but are not equivalent: the suppression index depends on the strength of surround influence for distractors matched to the target, whereas the pop-out index depends on the tuning of the surround (i.e., whether suppression is different for distractors matched to the target or different from it). We measured the association between these two indices using Pearson partial correlations, since both indices are calculated using responses to homogeneous displays. Partial correlations were significant in both V1 (0.77, p<0.001; Fig 3C) and V2 (0.64, p<0.001).

We wondered if the stronger suppression and pop-out modulation in V2 than V1 were due to the stimulus size and spacing in our displays, which were chosen with the larger V2 receptive fields in mind. To test this possibility, we ran control experiments in a subset of V1 units (n=10) with targets and distractors better matched in size to V1 preferences (2° in size and 0.6° spacing). With these smaller stimuli, the pop-out index (0.12±0.05) and suppression (0.24±0.06) in V1 were similar to those observed in V2 units for the larger displays.

We conclude that the size and spacing of our displays are appropriate for revealing pop-out modulation in both V1 and V2, though the magnitude of this modulation is sensitive to stimulus size and spacing.

### Pop-out modulation for displays of textures and noise images

We next asked whether there was pop-out modulation in V1 and V2 for displays in which targets and distractors differed only in the presence or absence of higher-level statistics in texture stimuli. Sensitivity to higher-level texture statistics is more evident in V2 than V1 (Freeman and Simoncelli, 2013; Ziemba et al., 2016, 2018), so we might expect pop-out modulation for these displays to be more evident in V2. If so, this would suggest that neural correlates of salience can exist independently of proposed V1 ‘salience maps’ (Li, 1999; Itti and Koch, 2001; Zhaoping and Zhe, 2015).

We synthesized naturalistic texture stimuli using the Portilla & Simoncelli algorithm (Portilla and Simoncelli, 2000). Each source texture image was the basis of a family, or a set of synthetic textures with the same set of summary statistics. To test the contribution of higher-level statistics—defined in the algorithm by the relationships among filter outputs applied to the image—we also synthesized control images for each texture family (termed ‘noise images’ hereafter). Noise images were matched to each synthesized texture family in spectral content, contrast, and luminance only. To ensure responses were not affected by a feature present within a particular instantiation of a synthesized texture, we used a different sample of each texture within the display on each trial. In our displays, the target could be either a noise image with texture distractors from the same family, or a texture target with noise distractors (Fig. 2C). We found no difference between these two cases for any of the metrics below, so the results are presented averaged across the two types of displays.

Most V1 and V2 units responded similarly to heterogeneous and homogeneous displays (Fig 4A, left); only a few units showed some response difference (Fig 4A, right). On average, responses were nearly identical for heterogeneous and homogeneous displays, as evident in the population average PSTHs for both V1 and V2 (Fig 4B).

**Fig 4.**
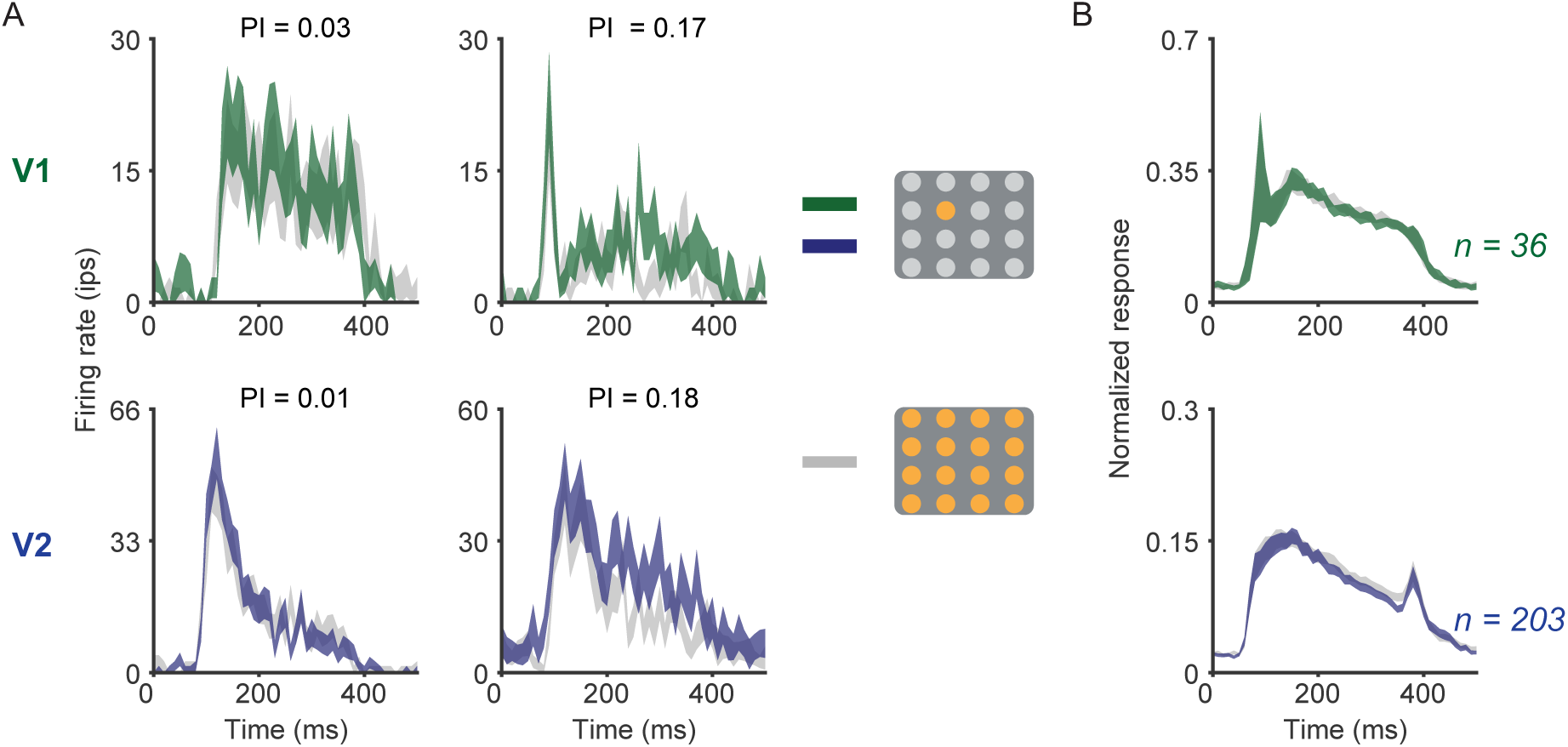
Responses to displays of textures and noise images. **(A)** Responses of example V1 (top) and V2 (bottom) units to heterogeneous (colored traces) and homogeneous (gray) displays. The thickness of the lines depicts S.E.M. across trials. Units in V1 and V2 usually showed little (left) but sometimes some (right) difference in responses to the two displays. Value of the pop-out index (PI) is indicated above each sublot. **(B)** Average PSTHs for V1 (top) and V2 (bottom) units. The PSTH for each cell was normalized by its peak value and these were then averaged across neurons.

To quantify differences in responsivity for heterogeneous and homogeneous displays, we calculated a pop-out index for each unit that had a significant response to the target only condition. Units in V1 had pop-out indices near zero (-0.005±0.009; p=0.6 for difference from 0, t-test; Fig 5A, top), indicating nearly identical responses to heterogeneous and homogeneous displays. V2 units had a slightly greater response for homogeneous than heterogeneous displays (-0.03±0.004; p<0.001, t-test; Fig 5A, bottom). Only a small percentage of units in each area showed significant pop-out modulation (V1=4.2%, V2=5.8%; bootstrap test), a percentage expected from the false positive rate given our criterion for statistical significance.

**Fig 5.**
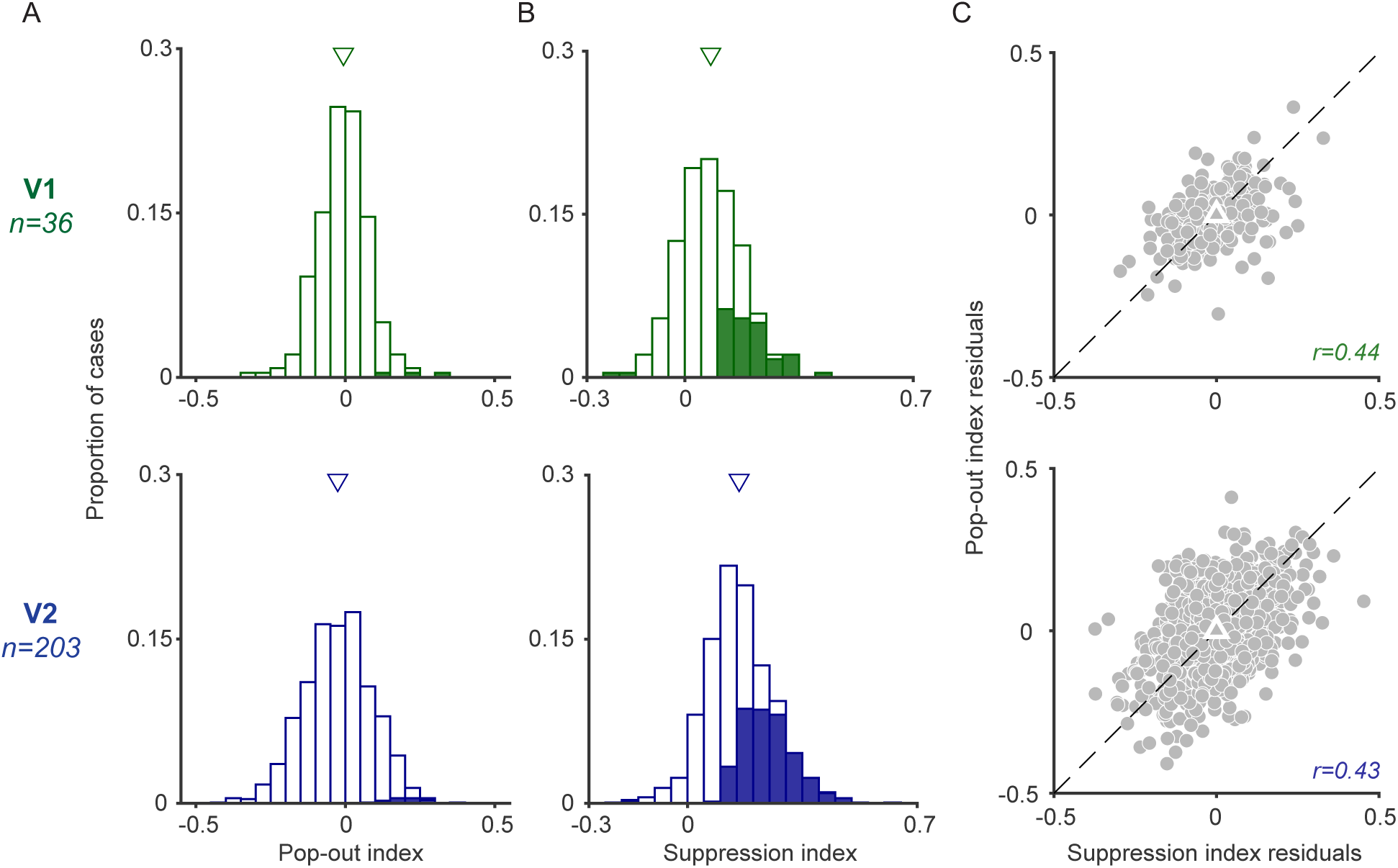
Pop-out modulation for displays of texture and noise. **(A)** Pop-out indices for V1 (top) and V2 (bottom) units. Shaded bars represent significant cases. Arrowhead indicates the mean value. **(B)** Suppression indices for V1 (top) and V2 (bottom) units. Shaded bars represent significant cases. **(C)** Relationship between suppression (abscissa) and pop-out (ordinate) index residuals, for V1 (top) and V2 (bottom).

To test whether the weak pop-out modulation was due to a lack of surround modulation from distractors, we computed suppression indices using responses to the target alone and homogeneous displays (Fig 5B). In both V1 and V2, distractors reduced responses to targets (suppression index of 0.11±0.01, p<0.001; and 0.17±0.004, p<0.001, respectively); suppression was stronger in V2 than V1 (p<0.001, unpaired t-test). A substantial percentage of V1 and V2 units were significantly modulated by the distractors (21.3% and 37%, respectively; bootstrap analysis). On average, suppression indices were similar for displays of gratings and of textures, for both V1 (p=0.86, unmatched t-test) and V2 units (p=0.57, unmatched t-test). The suppression and pop-out indices for displays of texture and noise were significantly correlated in V1 (partial correlation of 0.44, p<0.001; Fig 5C) and V2 (0.43, p<0.001).

Our results thus indicate that although noise distractors induce significant suppression of responses to texture targets (and vice versa), there is little pop-out modulation for these displays in either V1 or V2.

### Pop-out responses for displays with distinct texture families as targets and distractors

We next tested whether displays consisting of targets and distractors from different texture families evoke pop-out responses in V1 and V2. Unlike stimuli in the preceding experiment, the targets and distractors in these displays would have different spectral content as well as higher-order statistics. These displays thus allowed us to test if pop-out responses occur for stimuli which differ in both low-level (i.e. spectral) and high-order texture statistics. Though prior work has shown pop-out modulation for targets and distractors that differ in low-level statistics (e.g. orientation), it is not clear whether such modulation would occur when those low-level differences are embedded in more complex images.

The pop-out indices for these displays were small in both V1 (0.05±0.02; p=0.007, t-test) and V2 (0±0.01; p=0.99, t-test; Fig 6A). V1 units had slightly higher pop-out indices than V2 (p=0.01, unpaired t-test). Only a small percentage of units in each area had significant pop-out indices (12.1% in V1, 3.6% in V2; bootstrap analysis). We found weak but statistically significant surround modulation in V1 (0.09±0.02; p<0.001, t-test; Fig. 6B) and V2 (0.11±0.01; p<0.001, t-test; p=0.4, unpaired t-test for comparison of the two areas), though the suppression indices were smaller than those for grating and texture vs. noise displays. A small percentage of units had significant suppression indices in V1 (10.3%) and V2 (11.8%). The pop-out and suppression indices were correlated across units (in V1, partial correlation of 0.55, p<0.001; in V2, 0.47, p<0.001; Fig 6C). Units with stronger surround suppression thus had stronger pop-out modulation.

**Fig 6.**
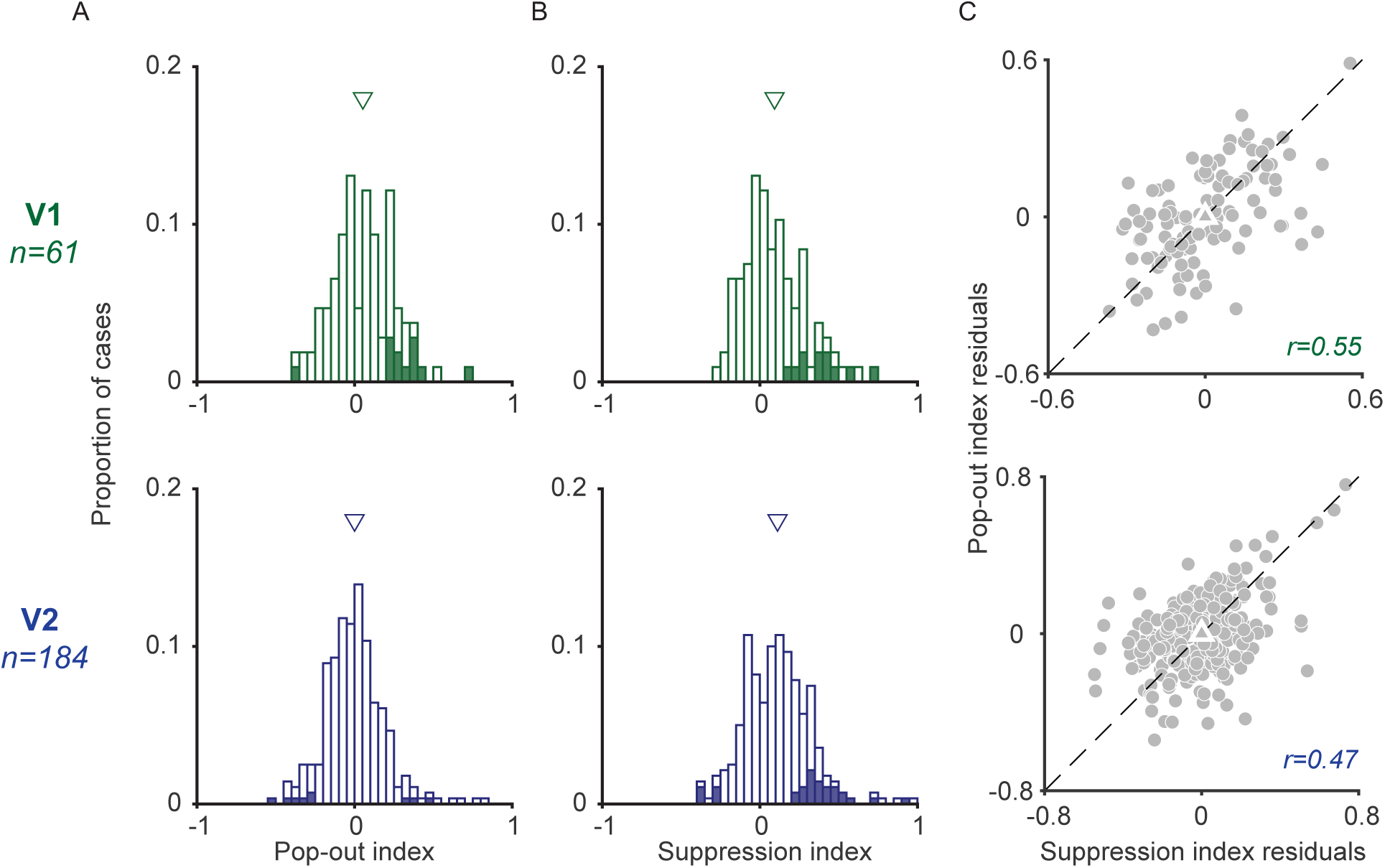
Pop-out modulation and suppression displays with distinct texture families as targets and distractors. **(A)** Pop-out indices for V1 (top) and V2 (bottom) units. Shaded bars represent significant cases. Arrowhead indicates mean. **(B)** Suppression indices for V1 (top) and V2 (bottom) units. Shaded bars represent significant cases. **(C)** Relationship between suppression (abscissa) and pop-out (ordinate) index residuals for V1 (top) and V2 units (bottom).

We conclude that displays in which a target and distractors are from different texture families also fail to evoke pop-out responses, in either V1 or V2.

### Effects of adaptation on pop-out responses for textures

Previous perceptual (McDermott et al., 2010; Wissig et al., 2013) and physiological (Dutta et al., 2016) work has provided evidence that adaptation can increase salience and pop-out modulation.

We tested whether adaptation could also modulate, or induce, pop-out responses for texture displays.

To examine the effects of adaptation on pop-out modulation, we measured the pop-out index after presentation of a 600 ms adapter (Fig 7A), a duration sufficient to induce adaptation effects in V1 and V2 units (Davila and Kohn, 2023). The adapter on each trial (Fig 7B) was either: (1) a homogeneous display of textures, (2) a homogeneous display of noise images, or (3) a blank screen (used to measure control or pre-adaptation responses). The type of adapter was chosen randomly on each trial. Each image patch in each display contained a unique sample, at each location and on each trial.

**Fig 7.**
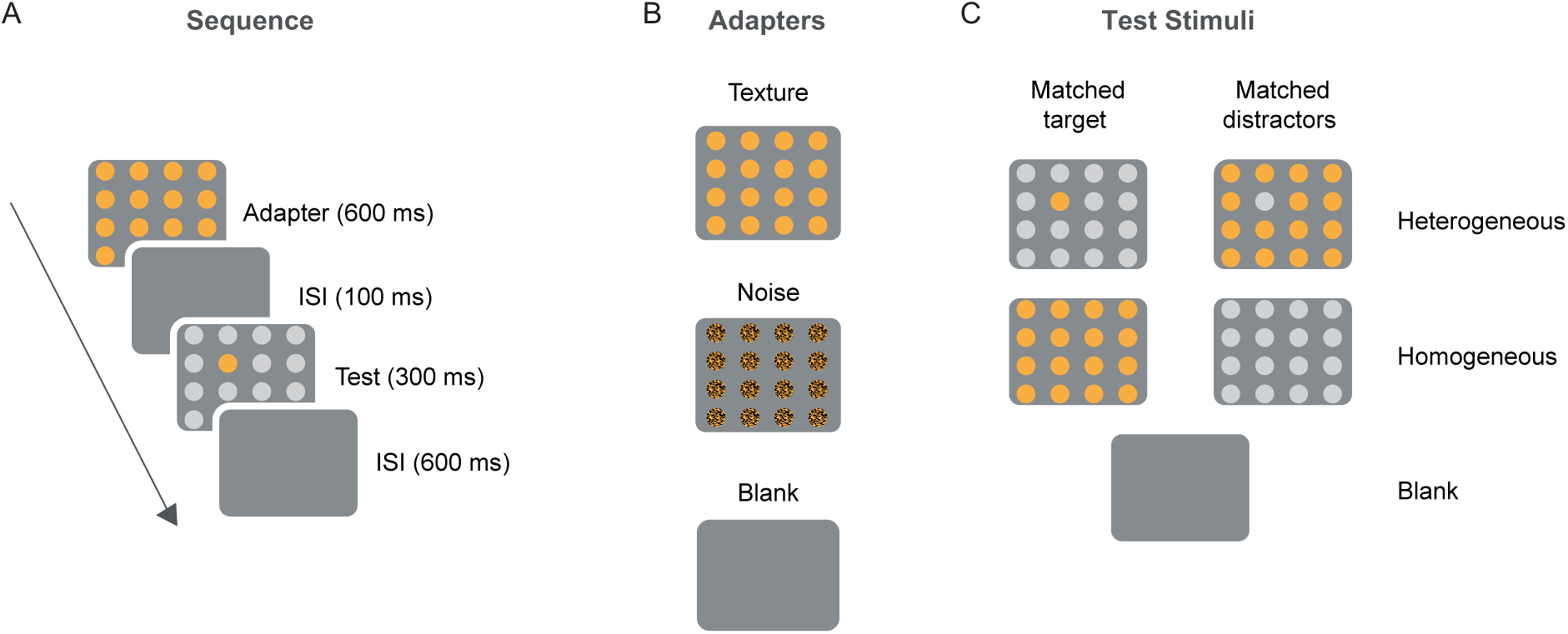
Design of adaptation experiments. **(A)** Each trial consisted of an adapter followed by a blank and a test stimulus. **(B)** Adapters consisted of either a grid of textures, noise images, or a blank. Texture and noise grids consisted of images from the same texture family, displayed at locations matched to the target and distractors of the test stimuli. **(C)** Test stimuli were heterogeneous, homogenous, target alone or distractors alone displays. Additionally, the adapter family could be matched to either the targets (left) or the distractors (right) of the test stimulus

Test stimuli consisted of targets and distractors from different texture families. We measured responses to displays in which (1) the target was matched to the texture family of the adapter (Fig 7C, left column) or (2) the distractors of the heterogeneous display were matched to that adapter (Fig 7C, right column). We also included in our test ensemble each target on its own and the two sets of distractors alone (not shown). These stimuli were used to select the neurons which were responsive to the target but not driven by the distractors. Displays with targets alone were also used in computing the suppression index across units.

We focused our analyses on the measured V2 responses. This is because V2 seemed a more promising candidate area for inducing pop-out modulation for textures, since responses there are modulated by both low-level and higher-order texture statistics (Freeman et al., 2013). Additionally, adaptation effects for textures are more stimulus specific in V2 than in V1 (Davila and Kohn, 2023). We did record a smaller number of V1 units in these experiments as well, but the pre-adaptation pop-out indices were different for displays in which either the target or the distractors were matched to the adapter (left vs. right column of Fig 7C; 0.14±0.02 vs. - 0.03±0.02, p <0.0001). We ascribe this difference in pop-out responses to the relatively small sample size (matched target: *n*=52, matched distractors: *n*=55) and were thus not confident in drawing conclusions about the effects of adaptation in these data.

We first measured how V2 responses changed after adaptation when the adapter was matched to the test target. Before adaptation (i.e., after presentation of a blank adapter), the pop-out index was near zero (-0.09±0.01, p=0.2, t-test; Fig 8A). After adaptation, the pop-out index increased to 0.11±0.02 (p<0.001, paired t-test; Fig 8A). To test if the increase in pop-out modulation was driven by the spectral content or the higher-level statistics of the adapter, we analyzed responses measured after presenting a noise adapter. This adapter also increased the pop-out index in V2 (0.11±0.02, p<0.001 matched t-test; Fig 8B); the increase in the pop-out index was statistically indistinguishable for texture and noise adapters (p=0.99, unmatched t-test). Thus, the increase in pop-out modulation after adaptation is likely due to effects induced by the spectral content of each texture, consistent with past evidence that adaptation effects in early visual areas are orientation and spatial frequency specific (Movshon and Lennie, 1979; Saul and Cynader, 1989; Müller et al., 1999; Dragoi et al., 2000; Crowder et al., 2006; Patterson et al., 2014a).

**Fig 8.**
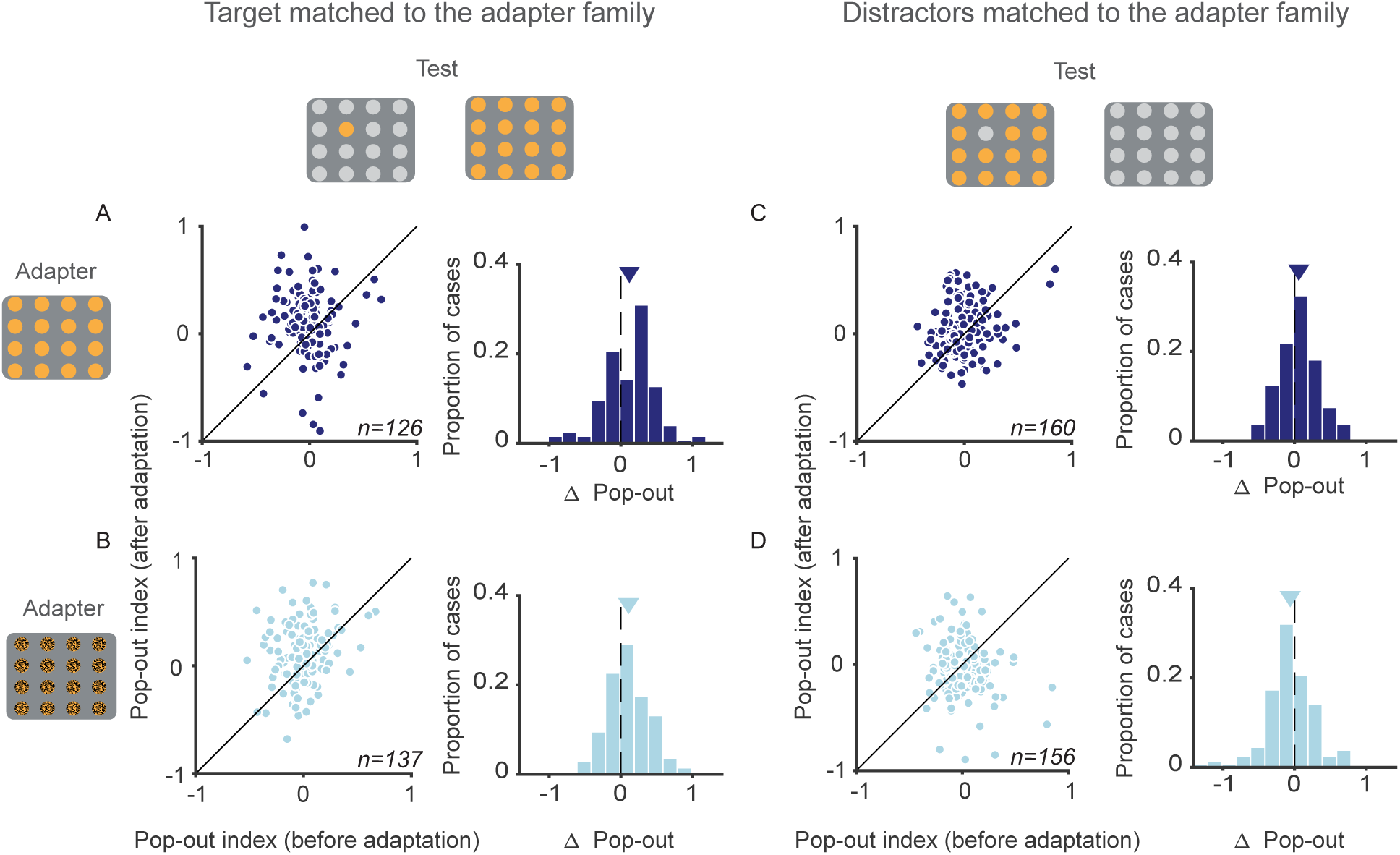
Effect of adaptation on pop-out modulation in V2. **(A)** Pop-out indices before (abscissa) and after (ordinate) adaptation, when the texture family used as an adapter is matched to the target. Histograms to the right indicate the change in pop-out index after adaptation (after adaptation compared to before adaptation). (**B**) Pop-out indices after adaptation with a noise stimulus matched to the target family. **(C)** Pop-out indices when the texture family used as an adapter is matched to the distractors of the heterogeneous display. **(D)** Pop-out indices after adaptation with a noise stimulus from the same texture family as the distractors of the heterogeneous display.

We next tested whether adapting V2 units to textures matched to the distractors of heterogeneous displays would also alter pop-out modulation. Based on previous work, adapters matched to the distractors should influence the suppression they provide and thus might affect pop-out modulation (Webb et al., 2005; Camp et al., 2009; Wissig et al., 2013; Patterson et al., 2014a). Before adaptation with these displays, the pop-out index was 0±0.02 (p=0.96, t-test; Fig 8C). After adaptation the pop-out index increased to 0.06±0.02 (p<0.01 for comparison to pre-adaptation values, matched t-test; Fig 8C). Adapting with noise images from the same family as the distractors had the opposite effect: it caused the pop-out index to decrease slightly (- 0.06±0.02; p=0.02, matched t-test; Fig 8D).

Thus, adapters matched to targets or distractors are both capable of altering pop-out modulation in V2. When the adapter is from the same family as the target of the test display, there is an increase in pop-out modulation. This increase is evident for both noise and texture adapters. When adapters are matched to distractors of the heterogeneous display, the effects are dependent on high-level statistics: texture adapters cause the pop-out index to increase, whereas noise adapters cause it to decrease. This difference could be due to sensitivity to texture statistics in the surround (Ziemba et al., 2018).

To understand more precisely how adaptation alters pop-out modulation, we examined the effects on responses to homogeneous and heterogenous displays separately—the two components that define the pop-out index.

We first analyzed responses to heterogeneous and homogenous displays containing a target from the same family as the adapter (Fig 9A). Since targets were matched to the adapter for both displays, differences in the adaptation effects between these displays would be due to adaptation specificity for the surrounding stimuli (i.e., distractors).

**Fig 9.**
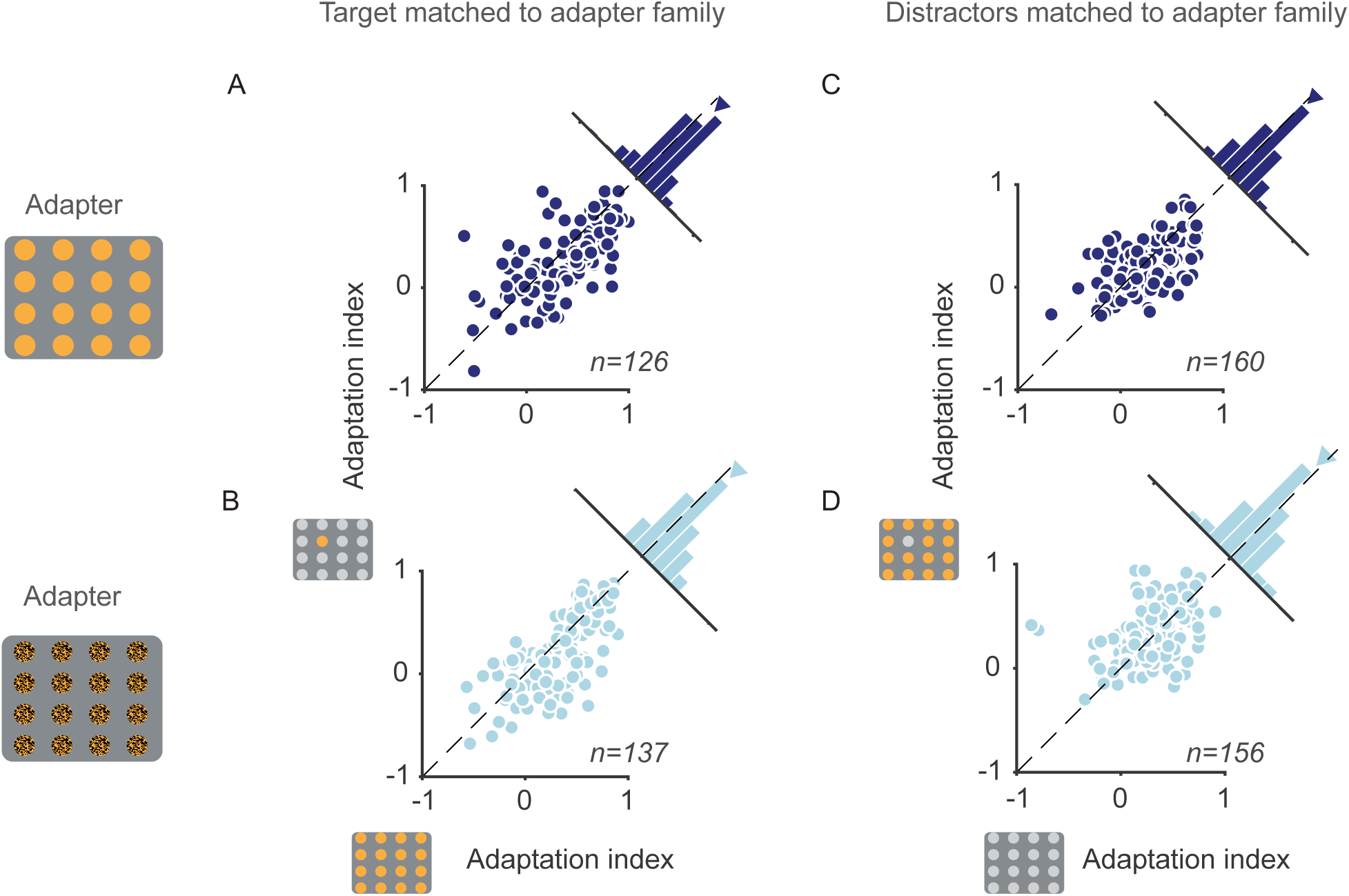
The effects of adaptation on the target and distractors. **(A)** V2 adaptation indices following a texture adapter (top grid, dark blue) matched to the target, for homogenous (abscissa) and heterogeneous displays (ordinate). **(B)** Adaptation indices following a noise adapter (bottom grid, light blue) matched to the target, for homogenous (abscissa) and heterogeneous displays (ordinate). **(C)** Adaptation indices for homogeneous (abscissa) and heterogeneous displays (ordinate) in which the target was different from the texture adapter; the distractors were matched to the adapter for the heterogeneous display. **(D)** Adaptation indices for homogenous (abscissa) and heterogenous displays (ordinate) in which the target was from a different family from the noise adapter. Mean adaptation indices are plotted with triangle symbols in all plots.

To quantify adaptation effects, we calculated an adaptation index defined as the difference in responsivity before and after adaptation divided by the sum. Adaptation indices were positive for homogenous (0.38±0.03, p<0.001, t-test; Fig 9A) and heterogeneous displays (0.29±0.03, p<0.001, t-test), indicating that adaptation caused a substantial loss of responsivity. The indices were larger for the homogeneous displays (p=0.002, matched t-test). Similarly, following a noise adapter from the same family as the target, adaptation indices were higher for homogeneous (0.35±0.03, p<0.0001, t-test) than heterogeneous conditions (0.25±0.03, p<0.0001, t-test; p=0.02 for comparison, matched t-test). Thus, the increase in the pop-out index after presentation of either texture or noise adapters arose because responses to homogeneous displays were reduced more than those to heterogeneous displays after adaptation.

We next examined the cases in which the adapter was matched to the distractors of the heterogeneous displays. In these data, adaptation indices were also higher for homogenous (0.27 ±0.02, p < 0.001, t-test; Fig 9B) than heterogeneous displays (0.22±0.02, p < 0.001, t-test; p=0.02 for comparison, matched t-test; Fig 9B). Higher adaptation indices for homogenous than heterogenous conditions cause an increase in the pop-out index after adaptation. On the other hand, noise adapters matched to distractors elicited similar adaptation indices for homogeneous (0.28±0.02, p<0.0001, t-test) and heterogeneous conditions (0.33±0.02, p<0.0001, t-test), which were not significantly different from each other (p=0.11, matched t-test). The stronger response reduction for heterogeneous displays explains the slight decrease in pop-out indices after adaptation with noise adapters matched to the distractors.

## DISCUSSION

We measured pop-out modulation in V1 and V2 to displays of texture patches. Our goals were to test 1) whether pop-out modulation could occur for features that were encoded after V1; and 2) if pop-out modulation could be altered by adaptation. Although there was robust pop-out modulation for displays of gratings, in both V1 and V2, there was little or no pop-out modulation evident for displays in which target and distractors were textures and noise images, or images from different texture families. However, adapting with homogeneous displays of textures or noise images could induce pop-out modulation for these displays in V2.

Our experiments—particularly those involving adaptation—required stable recordings for long periods, which we accomplished by recording in sufentanil anesthetized animals. While we cannot exclude the possibility that our results were influenced by anesthesia, the available evidence suggests that similar results would be obtained in awake animals. First, sufentanil has little effect on activity in early sensory cortex (Constantinople and Bruno, 2011). Second, adaptation effects are similar in awake and sufentanil-anesthetized primates (Müller et al., 1999; Dragoi et al., 2000, 2002; Patterson et al., 2013, 2014a), and texture selectivity develops gradually across cortical areas in a similar manner in sufentanil-anesthetized (Freeman et al., 2013; Ziemba et al., 2016, 2019, but see Ziemba et al., 2024) and awake primates (Okazawa et al., 2015, 2017). Third, robust pop-out modulation has been reported in V1 of anesthetized macaques (Nothdurft et al., 1999), cats (Kastner et al., 1997), and owls (Dutta et al., 2016).

Finally, surround suppression for gratings (Hubel and Wiesel, 1965; Bair et al., 2003; Cavanaugh et al., 2002b) and textures (Ziemba et al., 2018) is also robust under anesthesia in both V1 and V2.

### Pop-out along the visual hierarchy

Influential models have proposed that salience is defined by response differences among V1 neurons encoding stimuli at different spatial locations (Li, 1999, 2002). The differences in V1 responsivity to simple features (e.g. color, orientation, motion, size) across a visual field— influenced by surround suppression—can explain perceptual salience via a feature map, which is thought to be constructed in a bottom-up parallel manner and to drive attention automatically (Treisman and Gelade, 1980).

Though feature maps are often assumed to be represented in V1, some physiological studies have shown that V1 responses are inconsistent with perceptual salience: conjunction-based target stimuli are not perceptually salient (Treisman and Gelade, 1980; Treisman and Gormican, 1988; Wolfe, 1994), yet evoke robust pop-out modulation in V1 (Hegdé and Felleman, 2003). In V4, on the other hand, there is stronger pop-out modulation for salient singleton features than conjunctions, suggesting responses downstream of V1 may provide better correlates of salience (Burrows and Moore, 2009). Pop-out responses have also been reported in V2 (Lee et al., 2002), V4 (Mazer and Gallant, 2003; Burrows and Moore, 2009), lateral intraparietal area (Buschman and Miller, 2007), and the superior colliculus (White et al., 2017). Whether this modulation is computed in V1 and inherited by these areas or computed *de novo* in some or all of these areas is unclear.

To test whether neurons in higher visual areas support pop-out modulation independently of V1, we used displays of naturalistic textures. Since V1 neurons are insensitive to the high-level statistics of textures (Freeman et al., 2013), they should respond similarly to homogenous and heterogenous displays of naturalistic textures and noise, assuming similar insensitivity for texture statistics in the V1 surround. Since V2 responsivity does depend on these statistics (Okazawa et al., 2017; Ziemba et al., 2016, 2019), V2 units might respond differently to targets embedded in these homogeneous or heterogeneous displays. However, we observed no pop-out modulation for targets surrounded by noise distractors (or vice versa), in either V1 or V2. In contrast, displays of gratings, whose size and spacing were the same as for texture displays, resulted in clear pop-out modulation in both V1 and V2.

The absence of pop-out modulation for texture displays was not due to a lack of surround modulation: there was robust suppression of responses to targets by distractors in both areas. We note that because surround suppression decreases with retinotopic distance (Cavanaugh et al., 2002b; Bair et al., 2003), pop-out modulation and surround suppression will depend on target and distractor size and their spacing. Since we found higher pop-out indices for units whose responses were more strongly suppressed by distractors, other combinations of target size, distractor size or density, and patch spacing—all of which will alter the strength of surround suppression—might yield more robust pop-out modulation.

We also found little pop-out modulation in V1 and V2 for texture targets surrounded by distractors from other texture families. Different texture families not only have distinct high-level statistics, but also differ in their spectral content (Portilla and Simoncelli, 2000). Since V1 neurons are sensitive to image spectral content, displays of textures and distractors from different texture families might be expected to recruit robust pop-out modulation in V1. We did find slightly higher pop-out modulation for these displays in V1 than V2. The absence of more compelling pop-out modulation might be attributed to the broad spectral content of most texture images, and the broad tuning for spatial frequency in the surround (Webb et al., 2005; Wissig and Kohn, 2012). As a result, the differences in surround suppression provided by matched (homogeneous displays) and mismatched (heterogeneous displays) distractors might be small.

Our findings are consistent with perceptual work showing that higher-order texture statistics do not modulate salience. Julesz proposed that differences in texture statistics beyond second-order cannot be processed by bottom-up mechanisms and instead require serial top-down attention—the Julesz conjecture (Julesz, 1981). However, violations of this proposal have been reported, including by Julesz himself (Julesz, 1981; Victor et al., 2017). For instance, two textures with identical local second-order statistics can be easily (pre-attentively) discriminated if the constituent elements in one of the textures are arranged to have some degree of collinearity (Caelli et al., 1978). Such statistics are reminiscent of the correlations of filter outputs across space which is one of the defining higher-order statistics of the Portilla and Simoncelli (2000) textures we used. Other perceptual work has shown that complex features not defined by singleton features, such as shape from shading, can be salient (Ramachandran, 1988). Our results suggest that if texture targets amid either noise or other texture distractors are perceptually salient, the relevant signals would have to be computed downstream of V1 and V2.

### Adaptation and pop-out

One function of adaptation might be to modulate salience, determining in part which features in the environment are of interest (Kohn, 2007). Because adaptation alters how neurons are affected by stimuli both in the receptive field center and surround (Webb et al., 2005; Wissig and Kohn, 2012; Patterson et al., 2013), it can easily alter responses to targets and distractors and their interaction. Consistent with this suggestion, adaptation to distractors increases perceptual salience in a search task (McDermott et al., 2010; Wissig et al., 2013). In the owl’s optic tectum, pop-out modulation is altered by adaptation to target and distractors of different orientations (Dutta et al., 2016). Additionally, salience and pop-out modulation can increase after prolonged training with a particular display, indicating both short-term and long-term plasticity of these signals (Julesz, 1981; Treisman and Gormican, 1988; Lee et al., 2002; Yan et al., 2018).

We found that adaptation with displays of textures, matched to either targets or distractors, could induce pop-out modulation in V2. When the adapter was matched to the target texture, effects were similar for texture and noise adapters, indicating that the low-level image statistics are sufficient to alter pop-out modulation. The increase in pop-out modulation was due to a stronger reduction of responsivity for homogenous than heterogenous displays following adaptation, a surprising outcome. Homogeneous and heterogeneous displays contain the same target stimulus, which is matched to the adapter; so, responses to the target stimulus should be reduced similarly in the two cases. Distractors in the homogeneous display are matched to the adapter, so the suppression they provide should be weakened by adaptation (Webb et al., 2005; Camp et al., 2009; Wissig and Kohn, 2012; Patterson et al., 2013) partially offsetting the loss of responsivity to the target. If adaptation effects for distractors are specific to texture identity as prior work suggests they should be (Davila and Kohn, 2023), surround suppression should be better maintained for heterogeneous displays, resulting in a greater loss of responsivity than for homogeneous displays.

The pattern of results we observe—opposite to that expected from the reasoning above— could be explained by strengthened surround suppression after adaptation. Westrick et al. (2016) proposed that normalization signals, a concept encompassing multiple types of suppressive signals including those responsible for surround suppression, can be strengthened when the neurons providing and receiving suppression are consistently co-activated. This co-activation would be expected to occur with our adapters, which consistently pair a target stimulus with a set of distractors. Our prior work on normalization signals within receptive field (Aschner et al., 2018) lend credence to this theory. However, some effects of adaptation on V1 surround suppression for grating stimuli are seemingly consistent with the theory and others are not (Webb et al., 2005; Wissig and Kohn, 2012; Patterson et al., 2013; see Aschner et al., 2018 for discussion). Perhaps strengthened surround suppression occurs with adaptation in V2, or for more complex images, or because of details of the display or adaptation procedure.

Adapting with texture displays matched to the distractors also resulted in an increase in pop-out modulation, an effect not evident for noise adapters. The absence of a change in pop-out modulation might be due to specificity for texture statistics in the surround; that is, differences in V2 surround suppression between textures and noise stimuli (Ziemba et al., 2018).

Though the mechanisms responsible for altered pop-out modulation after adaptation remain uncertain, our results clearly indicate that pop-out modulation can be altered by recent visual experience. This finding, together with work of prior studies, suggests that a primary function of visual adaptation is to re-define which stimuli in the environment are most salient, based on recent visual experience.

